# Microplastic-mediated transport of PCBs? A depuration study with *Daphnia magna*

**DOI:** 10.1101/428151

**Authors:** Zandra Gerdes, Martin Ogonowski, Inna Nybom, Caroline Ek, Margaretha Adolfsson-Erici, Andreas Barth, Elena Gorokhova

**Author notes:** Corresponding author: Zandra Gerdes.

## Abstract

The role of microplastic (MP) as a carrier of persistent organic pollutants (POPs) to aquatic organisms has been a topic of debate. However, theoretically, the reverse POP transport can occur at higher relative contaminant concentrations in the organism than in the microplastic. The effect of microplastic on the PCB removal in planktonic animals was evaluated using the cladoceran *Daphnia magna* with a high body burden of polychlorinated biphenyls (PCB 18, 40, 128 and 209) exposed to a mixture of microplastic and algae (with 77% MP by mass); daphnids exposed to only algae served as the control. As the endpoints, we used PCB body burden, growth, fecundity and elemental composition (%C and %N) of the daphnids. We found that PCB 209 was removed more efficiently in the daphnids fed with microplastic, while there was no difference for the ΣPCBs between the microplastic-exposed and control animals. Effects of the microplastic exposure on fecundity were of low biological significance, even though both the starting PCB body burden and the microplastic exposure concentrations were high and greatly exceeding environmentally relevant concentrations.

## Introduction

Microplastic (MP, particles < 1 mm [1]) are emerging contaminant in our environments. Concerns have been expressed that microplastic may compromise feeding of aquatic organisms and facilitate transfer of organic pollutants in food webs [2,3]. Indeed, these small plastic fragments are being ingested by a variety of organisms, e.g. fish [4], bivalves [5], polychaetes [6], and zooplankton [7], with still unknown consequences. Commonly reported effects of microplastic exposure include decreased food intake [8] and increased chemical exposure, for example, via leakage of potentially toxic additives [9] or chemicals sorbed to microplastic particles surface from ambient water [10].

Filter-feeders are vulnerable to potential impacts of microplastics. Negative effects of pristine microplastic on food intake and growth have been reported for a range of zooplankton, e.g. the cladoceran *Daphnia magna* [11] and the copepod *Calanus helgolandicus* [8], albeit at high concentrations of microplastics used in the experiments. Moreover, for these animals at the lower trophic levels, the exposure to environmental contaminants absorbed to microplastic is of particular relevance for bioaccumulation in the food web and contaminant transfer to higher consumers. Therefore, it is important to establish whether organic and inorganic contaminants can be transferred when planktonic filter-feeders ingest microplastic [13,14]. Because persistent organic pollutants (POPs), such as polychlorinated biphenyls (PCBs), are hydrophobic, they have strong partitioning towards plastic and compartments that are rich in organic carbon, e.g. biota and sediment [14]. Due to their physicochemical properties, e.g., high plastic-water partition coefficient combined with a high surface to volume ratio, microplastic particles are efficient in sorbing hydrophobic substances [15]. Hence, microplastic can act as a vector for POPs between a consumer and its environment.

Several experimental studies that address microplastic-mediated POP transfer, from the environment to biota, were done by investigating non-contaminated animals exposed to microplastics loaded with POPs [10,16–18]. Such settings are, however, rarely relevant [19,20]. The ecologically plausible scenario should include consumers, microplastic and other compartments, such as natural suspensions of particulate matter, food, or sediment. Besseling et al. (2017) explored such a scenario by following PCB bioaccumulation in the lugworm *Arenicola marina*, with or without polyethylene microplastic added to the sediment [20]. In line with previous modelling studies [21,22], a limited microplastic effect was found – indicating that microplastic has limited relevance to the bioaccumulation *in situ*, as the ultimate direction of POP transport between the biota and plastic is a function of the the chemical fugacity gradient. In other words, if the POP concentration in the animal is higher than that in the microplastic, the contaminants will eventually be sorbed by the plastic and removed from the animal [23]. While this scenario is established in theory [22,23], the empirical data are still scarce. More experimental studies are, therefore, needed to validate models [19] and to understand the role of microplastic in POP transport.

We explored whether exposure to microplastics (1) facilitates depuration of PCBs, similar to the sought remediation effect by adding activated carbon to contaminated sediments [24], and (2) alliviates effects of the PCB exposure on growth (somatic and reproductive) and elemental composition (carbon, %C, and nitrogen, %N) in the model filter-feeder *D. magna* (Straus, 1820). These effects were studied experimentally by – first – feeding *D. magna* neonates with PCB-contaminated food to allow for PCB accumulation in the body. Then, the animals were offered a non-contaminated food with or without microplastic addition. The changes in the somatic and reproductive growth, elemental composition and PCB concentrations of the animals were evaluated at the end of the PCB exposure (juveniles) and the end of the depuration period (egg-bearing adults). We hypothesised that (I) the addition of microplastic to the system would facilitate removal of PCB from the daphnids, and, thus, a lower PCB body burden in the microplastic-exposed daphnids will be observed, and (II) a lower PCB body burden would result in higher somatic growth, fecundity and increased allocation of carbon and nitrogen, in the animals exposed to microplastic after PCB-exposure.

## Methods

### Test organisms

The freshwater cladoceran *Daphnia magna* (environmental pollution test strain *Clone 5*, the Federal Environment Agency, Berlin, Germany), were cultured in M7 media [25] at a density of 1 ind. 100 mL^−1^ and fed a mixture of green algae *Pseudokirchneriella subcapitata* and *Scenedesmus spicatus*. The algae were cultured in the MBL medium under constant light (70 μE cm^−2^ s^−1^) and temperature (24° C). Algal concentrations were determined using a 10AU™ Field Fluorometer (Turner Designs, Sunnyvale, California, USA) and established calibration curves.

### Polychlorinated biphenyls (PCBs)

Four PCB congeners (AccuStandard Inc., New Haven CT) covering a range of hydrophobicity within the substance group (Table 1, PCB 18, PCB 40, PCB 128 and PCB 209) were solved in toluene (AnalaR NORMAPUR VWR Chemicals) using equal proportions of each congener (by mass). This working stock was mixed with freeze-dried *Spirulina* (*Arthrospira* spp., Renée Voltaire AB, Sweden) and dried for 16 h under a stream of nitrogen to remove the toluene. The resulting absorption of PCB by *Spirulina* was quantified by gas chromatography-mass spectrometry (GC/MS, Table 1). The PCB-loaded *Spirulina* was suspended in 100 mL M7 and kept at +4° C before use in the feeding mixture.

**Table 1.** Log K_ow_ values and PCB concentrations in feed and *Daphnia*.

|  | Treatment | PCB 18 | PCB 40 | PCB 128 | PCB 209 |
| --- | --- | --- | --- | --- | --- |
| Log $K_{ow}$<sup>a</sup> | | 5.24 | 5.66 | 6.74 | 8.18 |
| *Spirulina*<sup>b</sup> |  | 410 (98) | 566 (75) | 441 (80) | 354 (69) |
| Accumulation phase<sup>b*</sup> | PCB | 7.66 (0.83) | 15.35 (1.53) | 7.60 (1.11) | 0.46 (0.06) |
| Depuration phase<sup>b*</sup> | PCB | 0.54 (0.15) | 1.40 (0.45) | 1.37 (0.43) | 0.37 (0.10) |
|  | PCB+MP | 0.47 (0.11) | 1.37 (0.26) | 1.50 (0.24) | 0.13 (0.04) |

<sup>a</sup> Log $K_{ow}$ values for PCB 18, 40 128 and 209 [30]. <sup>b</sup> congener-specific PCB content, $\mu\text{g}$ PCB g DW<sup>-1</sup>, in *Spirulina* (used as feed) and *Daphnia*<sup>\*</sup>. Values are shown as mean and SD.

## Microplastic

### Preparation of microplastic suspension

The microplastics (fluorescent green microspheres, FMG-1.3 1-5 μm, a proprietary polymer with density 1.3 g cm^−3^ and a melting point of 290° C) was purchased from Cospheric LLC (Goleta, USA). The concentration and particle size distribution of microplastics suspended in ultrapure water were measured with a laser particle counter PC-2000 (Spectrex, Redwood City, USA). The size range was confirmed to be 1-5 µm with a mean equivalent spherical diameter of 4±1.0 µm. To prevent adherence to the surface film and homo-aggregation of microplastic, a surfactant was added at a non-toxic concentration (0.01% w/w, Tween 80, P1754 Sigma-Aldrich)[26].

## Effects of PCB and microplastic exposure on growth and elemental composition

### Experimental outline

A two-step experimental procedure was used: (1) *accumulation phase* (day 0 to 3); the newborn daphnids were exposed to the PCB mixture via a contaminated food source, and (2) *depuration phase* (day 4 to 7); the juvenile daphnids were switched to a non-contaminated food with or without the addition of microplastic. At the termination of each step, daphnids were sampled to measure their body size as dry weight (DW) and body length (from the centre of the eye to the base of the apical spine, BL), elemental composition (%C and %N in the body) and PCB body burden (congener-specific).

At the start of the accumulation phase, *Daphnia* (< 24 h) in groups of ten were placed in glass beakers with 500 mL M7 media (n_total_ = 71, of which 40 replicates received PCB-loaded *Spirulina*). The experiment was conducted at 22°C with a 16:8 h light: dark cycle. After three days, 12 of the control and 14 of the PCB-exposed beakers were terminated to determine the size, elemental composition and PCB body burden at the end of the accumulation phase. The remaining beakers were divided into four treatments applied during the depuration phase: (1) *no-PCB control* with non-contaminated animals fed 100% algae (*control*, n = 11), (2) *microplastic control* with non-contaminated animals fed with the mixture of microplastic and algae, (*control*+*MP*, n = 8), (3) PCB-loaded daphnids fed 100% algae (*PCB*, n = 13), and (4) PCB-loaded daphnids fed with the mixture of microplastic and algae (*PCB*+*MP*, n = 13). The depuration phase lasted four days. Upon termination of either accumulation or depuration phase, the animals were transferred to 50 mL M7 media with 11 μgC mL^−1^ *P. subcapitata* for two hours to purge their gut contents [27] and minimize impact of the chemicals originated from the bolus. The daphnids were then assigned to samples designated for either DW and elemental composition or the BL and PCB analyses.

To standardise conditions and keep particle ingestion rates constant during the experiment, the daphnids were fed *ad libitum* [28] with a food mixture of *P. subcapitata* and *Spirulina*, provided at 1000:1 by dry mass. To maintain the saturating food levels, the media containing algae (8 μg C mL^−1^) and microplastic was renewed on days 0, 3 and 5. During the accumulation phase, the food mixture for the PCB-exposed animals contained PCB-loaded *Spirulina*, whereas the same amount of the non-contaminated *Spirulina* was used in the control. The PCB-loaded *Spirulina* powder was added to reach the exposure concentration of 3.20 μg PCB_tot_ L^−1^ (0.74 μg L^−1^of PCB 18, 1.02 μg L^−1^ of PCB 40, 0.80 μg L^−1^ of PCB 128, and 0.64 μg L^−1^ of PCB 209). These target concentrations were determined in a 48 h acute toxicity test with juvenile *D. magna* [25] that was conducted to establish the concentration causing < 10% mortality and measurable PCB body burden (S1 Text). The microplastic concentration during the depuration phase was 7 × 10^5^ particles mL^−1^, which equalled the algal concentration. By mass, microplastics and algae contributed approximately 77% and 23% to the total suspended solids in the system, respectively. According to a mass balance (S2 Text), this amount of microplastic was expected to produce a measurable difference in the *Daphnia* body burden between the treatments at the end of the depuration phase.

### Elemental analysis

The animals designated for the DW and elemental analyses were placed in pre-weighed tin capsules (4-10 individuals sample^−1^ depending on body size, n = 4-5 samples treatment^−1^), dried at 60° C for 24 h, weighed using a Sartorius M5P electronic microbalance, and stored in a desiccator. The C and N content was expressed as a mass fraction of DW (%C and %N, respectively). The analytical precision of was 0.9% for %C and 0.2% for %N (S4 Text).

### PCB analysis

All daphnids used for the PCB analysis were scanned using CanoScan 8800F flatbed scanner with ArcSoft PhotoStudio 5.5, and the images were used to determine BL (mm) and fecundity (embryos female^−1^). Using a weight-length regression established for *D. magna* (Eq. 1) [29], the individual DW was estimated as:

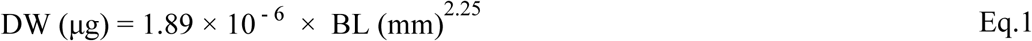

For the PCB analysis, daphnids were pooled by replicate (alive individuals, ~10 sample^−1^), homogenised with bead beating (acid-washed glass beads, 212-300 μm, Sigma-Aldrich) using FastPrep (MP Biomedicals), and stored in borosilicate test tubes (-18° C) until extraction. Internal standards were added (^13^C-PCB 128 and ^13^C-PCB 209) and the samples were extracted by shaking and sonicating twice with 5 mL n-hexane. The extracts were pooled, and 5 ml of concentrated sulphuric acid (H_2_SO_4_) was added to remove lipids. The samples were reduced in volume (200 µL), and volumetric standard (PCB 52) was added prior to instrumental analysis. The same PCB extraction and clean-up method was applied to PCB-loaded *Spirulina* samples. The samples (1 µL) were injected in split/splitless mode into a Thermo Scientific ISQ LR GC/MS equipped with a 30 m × 0.25 mm TG-SILMS column of 25 µm thickness. The column was held at 60° C for 2 min and then raised to 325° C at a rate of 30° C min^−1^. The congener mass per sample and individual DW were used to express PCB concentration; for quality assurance of the PCB analysis, see S3 Text.

### Data analysis and statistics

A one-compartment model was used to describe the elimination behaviour of PCB from a daphnid body:

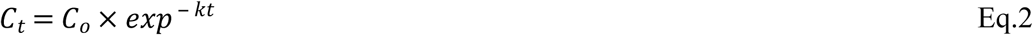

Where *C_t_* is the concentration of a PCB congener at the time (*t*) in the depuration phase and *C_o_* is the concentration at the onset of depuration (µg PCB g DW *Daphnia*^−1^). The elimination rate constant was determined by first estimating the overall elimination rate constant and then adjusting the rate constant by the growth rate constant for an individual, in order to account for growth dilution as a factor in determining the elimination rate kinetics:

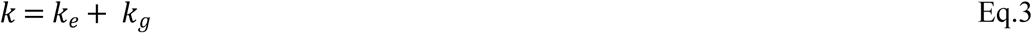

Where *k* is the overall elimination rate constant (d^−1^), *k_e_* and *k_g_* are the final first-order elimination rate constant (d^−1^) and the growth rate constant (d^−1^), respectively. For each replicate, *k_g_* was calculated as:

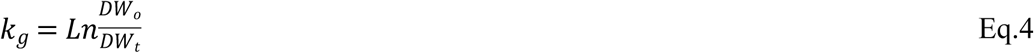

where *DW_o_* is the mean individual dry weight for a treatment group at the end of the accumulation phase, and *DW_t_* is the mean individual dry weight of a replicate at the end of the depuration phase.

Welch Two Sample t-test was used to compare PCB concentrations between the *PCB* and *MP*+*PCB* treatments. Generalized linear models (GLMs) with log link and normal error structure were used to evaluate the effects of the PCB and microplastics exposure on the response variables (DW, BL, %C, %N, fecundity and mortality). The PCB effects on DW, %C, and %N were evaluated in juveniles at the end of the accumulation phase and in the ovigerous females at the end of the depuration phase. In addition, the effects on the fecundity and body concentration of PCBs in the adults were evaluated for each PCB congener. The data were Box-Cox transformed, and standardised residual distributions and Shapiro-Wilks test for normality were used to assess the models. The most parsimonious models were selected based on the Akaike Information Criterion (AIC). Pairwise comparisons were tested using non-parametric Mann-Whitney tests. The significance level was set to α = 0.05 in all tests. S-plus 8.0 (TIBCO Software) and R version 3.2.3 were used for the statistical tests.

## Results

### Microplastic effects on the PCB elimination rate

At the end of the accumulation phase, the total PCB body burden had reached 30.4 ± 3.4 µg g *Daphnia*^−1^ (mean ± SD, Table 1 and Fig 1), and by the end of the depuration phase it had declined about 10-fold; no detectable levels were observed in any of the controls. The elimination rate of the PCB congeners decreased with increasing chlorination and decreasing polarity (Fig 2A). The mean *k_e_* values ranged from 0.55 d^−1^ for PCB 18 to 0.20 and 0.05 d^−1^ for PCB 209 in the PCB+MP and PCB treatments, respectively, with the corresponding half-lives of 1.0-2.0 and 4.1 d. A positive effect of microplastics on the elimination rate of PCB 209 was observed (Mann-Whitney, U = 0.5, p < 0.001, Fig 2A). Moreover, for this congener, the contribution of *k_e_* to the overall elimination rate constant *k* was significantly higher in the PCB+MP group (Mann-Whitney, U = 0, p < 0.001, Fig 2B). For the other congeners, *k_e_* contributed ~45-80% to their *k* values (Fig 2B), with higher values for congeners with lower logK_OW_ and no differences between the treatments.

**Fig 1.**
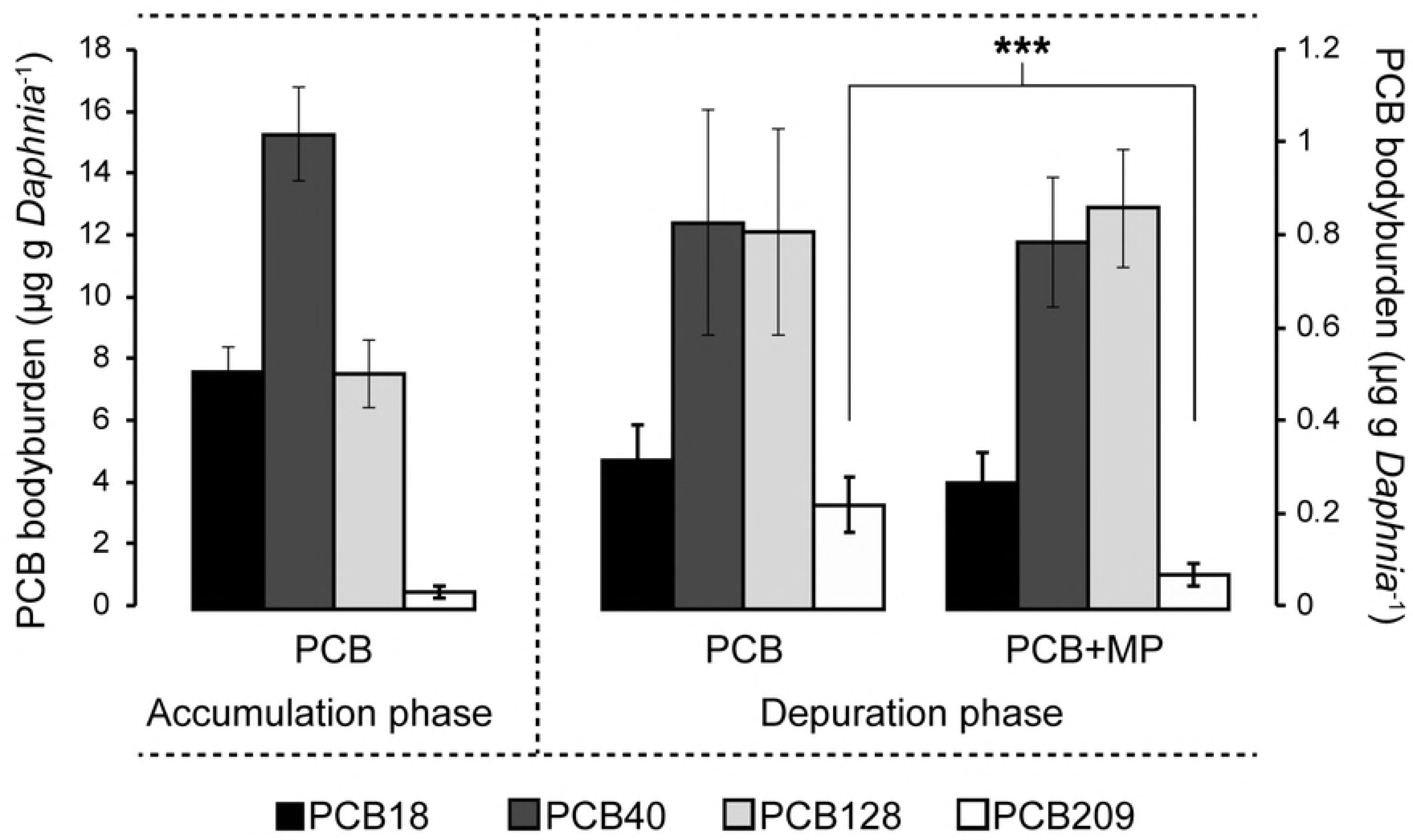
PCB Concentrations in *D. magna* before and after depuration. PCB concentrations (mean ± SD) in *D. magna* at the end of the accumulation (day 3, left y-axis), and depuration (day 7, right y-axis) phases; note the differently scaled y-axes. The concentration of PCB 209 was lower in the PCB+MP treatment compared to the PCB treatment (Welch Two Sample t-test, t_11_ = 5.6, p < 0.001). See also Table 1.

**Fig 2.**
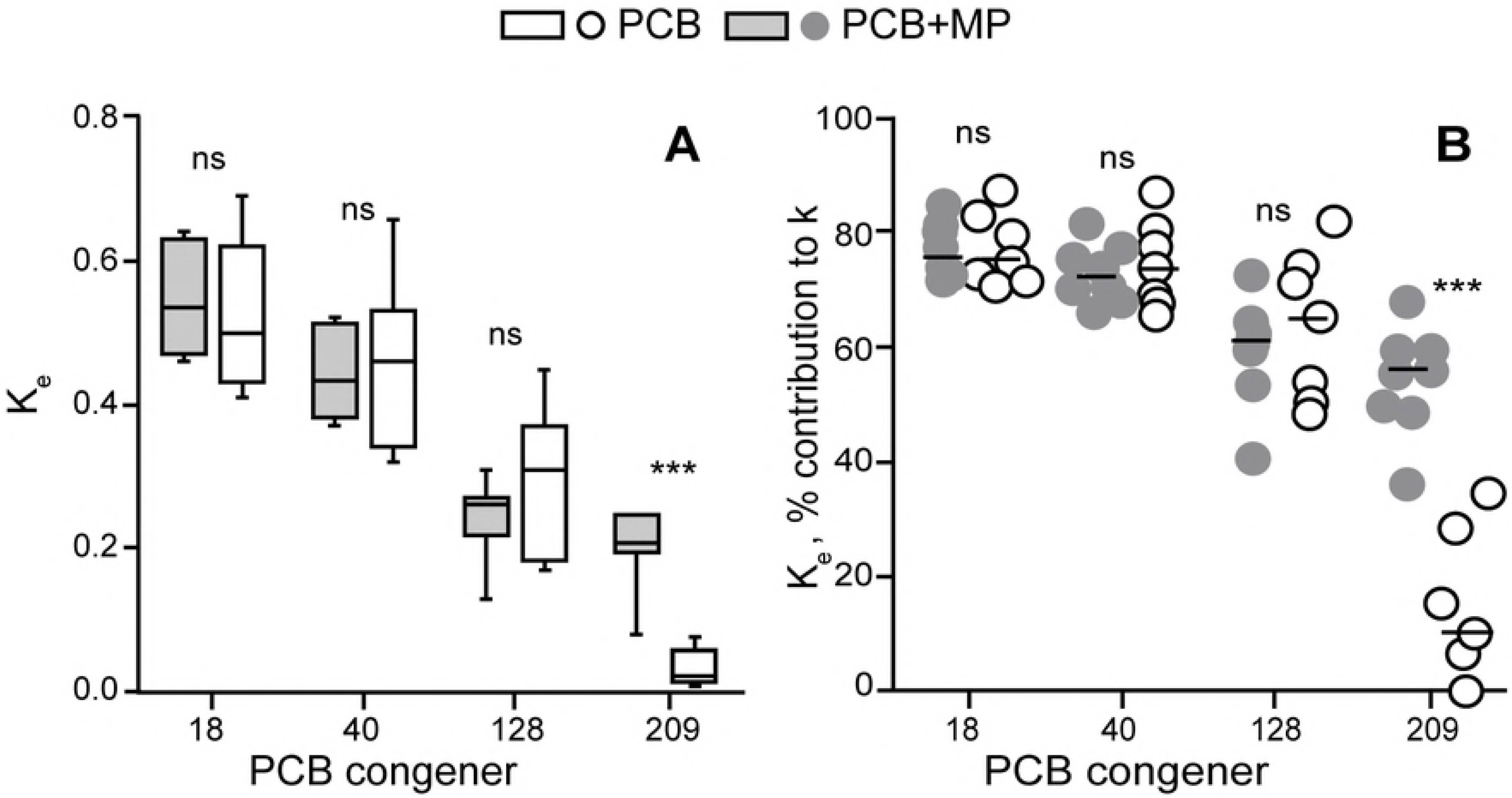
Elimination rate constants and the importance of growth dilution. Elimination rate constant for all PCB congeners (A) and contribution of k_e_ to the overall elimination rate (k) that accounts for growth dilution (B) during the depuration phase, ns: p > 0.05, ***: p < 0.001.

## Single and combined PCB and microplastic effects on the life history traits

### Mortality

In all treatments, mortality was low, not exceeding 7% during the accumulation phase and 6% during the depuration phase across the treatments. Exposure to microplastic did not affect mortality (GLM, Z_3,6_ = 0.10, p > 0.99 and Z_3,6_ = -0.10, p > 0.99 for the MP × PCB interaction and MP, respectively).

### PCB and microplastic effects

At the end of the accumulation phase, no significant effects of the PCB exposure on the body size, %C or %N in the juvenile daphnids were observed (Table 2, Fig 3A-C). At the end of the depuration phase, the PCB-exposed animals were significantly smaller (by 9% and 4% for DW and BL, respectively), whereas no microplastic effect on either DW or BL was observed (Table 2, Fig 3A). For all PCB-exposed daphnids, there was a significant negative relationship between the BL and PCB 209 concentration in the adults (GLM, Wald Stat._1, 11_ = 6.0, p = 0.01). Moreover, the %N values were significantly lower in the PCB-exposed daphnids compared to the control, with no additional effect exerted by microplastic (Table 2, Fig 3B). The %C values were also negatively affected by PCB exposure and, furthermore, a microplastic effect was observed, with %C values being significantly lower in the microplastic-exposed daphnids (Table 2, Fig 3C). The fecundity was positively related to the BL, with a significant positive effect of microplastic on the intercept of this relationship in the PCB-exposed animals (Fig 4, GLM, Wald Stat. _4, 9_ = 8.2, p < 0.01). This interaction implies higher size-specific egg production in the animals carrying PCB and receiving food mixed with microplastics. Fecundity in the PCB+MP treatment was also elevated in comparison to the control treatment receiving microplastic (control+MP, Fig 3D, GLM, t_1, 14_ = 3.3, p < 0.01).

**Table 2.** Summary of Generalized Linear Models results

| Phase | Dependent variable | Explanatory variable | Estimate | SE | Wald Stat. | p-value |
| --- | --- | --- | --- | --- | --- | --- |
| Accumulation | DW | PCB | -0.077 | 0.216 | 0.126 | 0.722 |
|  | %C | PCB | -0.005 | 0.007 | 0.400 | 0.508 |
|  | %N | PCB | -0.009 | 0.030 | 0.090 | 0.762 |
| Depuration | DW | MP | -0.005 | 0.023 | 0.053 | 0.818 |
|  |  | PCB | -0.060 | 0.023 | 6.721 | **0.009** |
|  |  | PCB * MP | -0.012 | 0.023 | 0.252 | 0.616 |
|  | BL | MP | -0.014 | 0.010 | 1.870 | 0.171 |
|  |  | PCB | -0.022 | 0.010 | 4.620 | **0.032** |
|  |  | PCB * MP | -0.007 | 0.010 | 0.420 | 0.518 |
|  | Fecundity | MP | -0.147 | 0.114 | 1.682 | 0.194 |
|  |  | PCB | 0.014 | 0.114 | 0.014 | 0.904 |
|  |  | PCB * MP | -0.280 | 0.114 | 6.063 | **0.014** |
|  | %C | MP | 0.000 | 0.000 | 4.700 | **0.029** |
|  |  | PCB | 0.000 | 0.000 | 6.700 | **0.010** |
|  |  | PCB * MP | 0.000 | 0.000 | 0.500 | 0.459 |
|  | %N | MP | 0.034 | 0.031 | 1.170 | 0.279 |
|  |  | PCB | -0.072 | 0.031 | 5.220 | **0.022** |
|  |  | PCB * MP | -0.009 | 0.031 | 0.080 | 0.078 |

GLMs testing treatment effects (PCB and MP against the control animals) and their interactions on the body size (DW, BL), fecundity, and elemental composition (%C and %N) in *D. magna* at the end of the accumulation and depuration phases. Significant effects are in bold.

**Fig 3.**
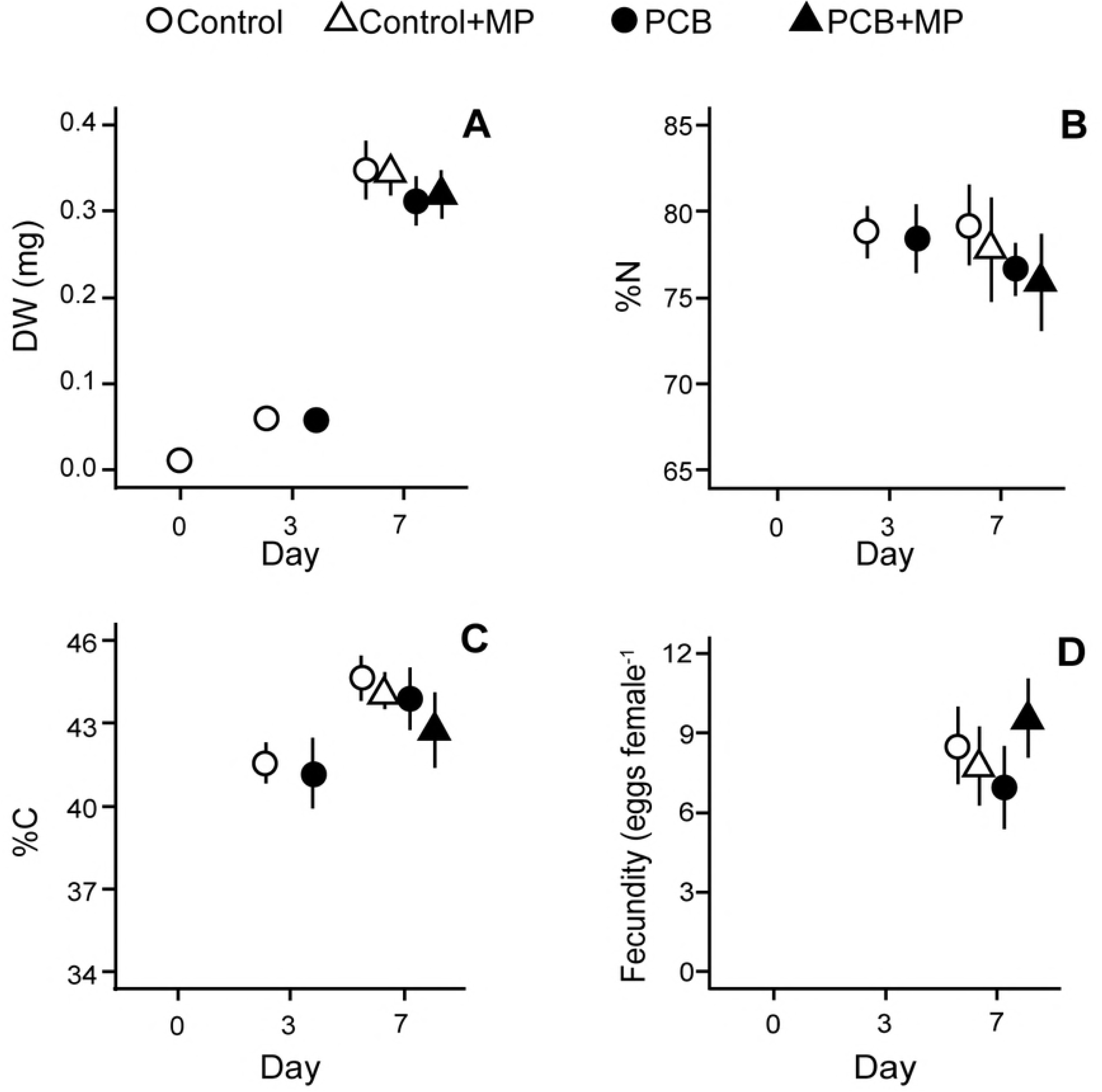
Treatment effects on physiological parameters of *D. magna*. The life-history parameters and body composition recorded on days 0, 3 (post accumulation phase) and 7 (post depuration phase) of the experiment: (A) individual body mass (DW), (B) carbon (%C) and (C) nitrogen content (%N), and (D) fecundity (eggs female^−1^). No eggs had been produced at the termination of the accumulation phase. Data are shown as mean ± SD. Some error bars are invisible due to the low SD.

**Fig 4.**
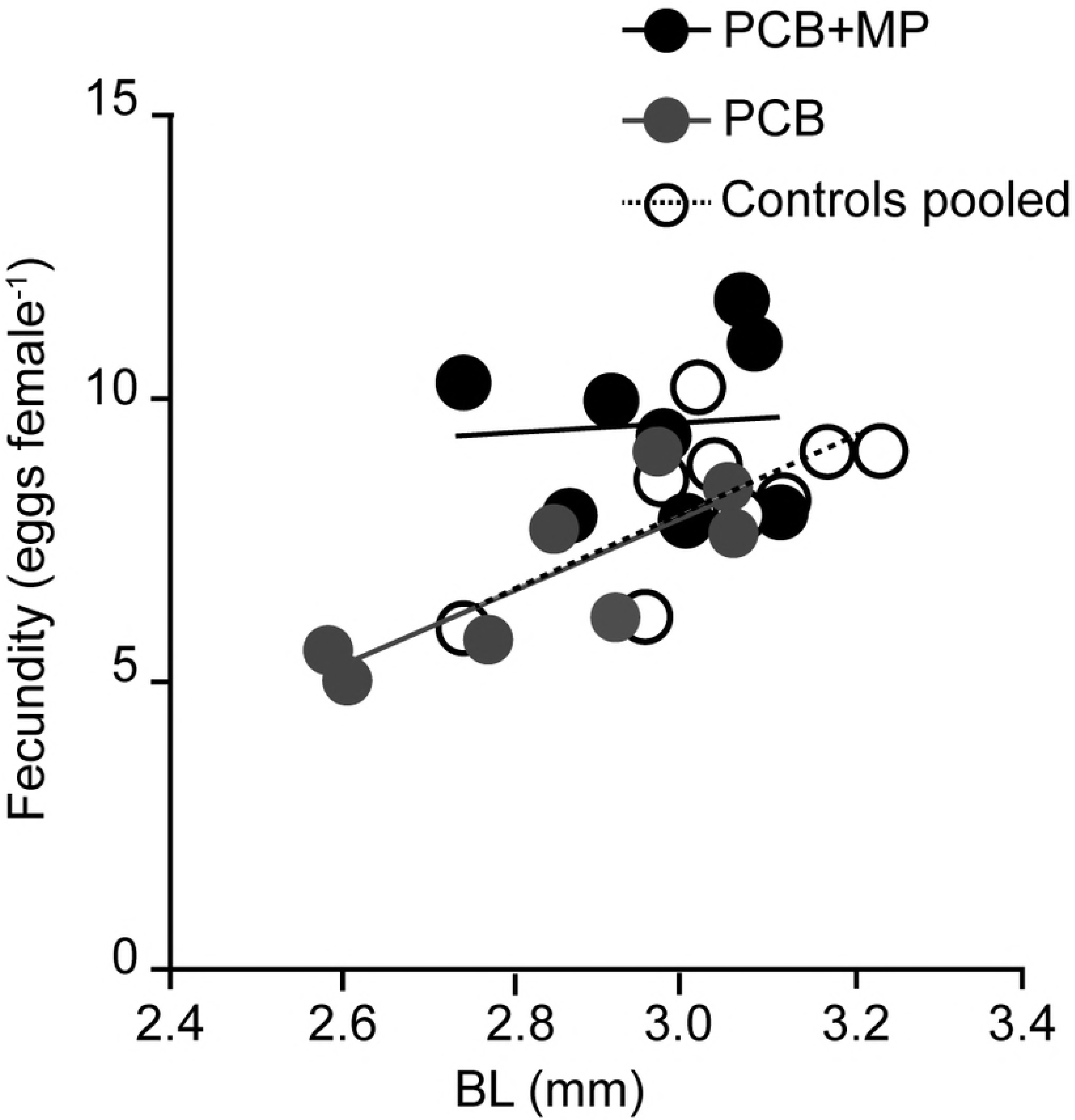
Treatment-specific relationships between fecundity and body length in *D. magna*. Whereas the overall relationship was significant, the group-specific slope was significantly different from zero only in the PCB group (R^2^ = 0.7, p < 0.02) and in the pooled controls (R^2^ = 0.5, p < 0.05).

## Discussion

The role of plastic debris as a carrier of POPs to aquatic biota has been much debated, but most of the evidence suggests that the bioaccumulation is mainly governed by the ingestion of natural prey [31]. Here, we tested whether microplastic can increase the transport of PCBs from small zooplankters with short gut residence time using a scenario that included both food and microplastic in the exposure system. According to the mass balance model of our depuration system (S2 Text), we anticipated a higher PCB 209 elimination in the daphnids receiving microplastic compared to those fed only algae. This was indeed the case, whereas there was no effect on the ΣPCBs despite the high plastic amount in the experiment. Thus, our first hypothesis that microplastics can increase PCB elimination in daphnids was confirmed, at least for the high-molecular-weight congeners.

The positive relationship between hydrophobicity of the PCBs and the contribution of growth to its elimination (Fig 2B) was expected, as more hydrophobic substances have longer half-lives [32]. The half-lives we observed (1-4 days) are reasonable to expect when considering the general importance of surface area over volume for chemical equilibration time. For instance, longer half-lives were reported for PCBs in larger crustaceans, e.g. 11 d for PCB 153 in the shrimp *Palaemonetes varians* [33] and 30 d for a PCB-mixture (PCB 40-81) in the amphipod *Diporeia spp*. [34]. In the copepod *Acartia tonsa* with a body size slightly smaller than that in *D. magna*, half-lives of 0.5-1 d for Aroclor 1254 (mainly penta and hexa-chlorinated biphenyls [35]) were reported [36].

The slower elimination of PCB 209 compared to the other congeners was most pronounced in the microplastic-free depuration treatment, resulting in a half-life of 4.1 d, whereas addition of microplastic decreased it to 2.0 d (Fig 2A). This difference is interesting and it shows that our microplastic were effective sorbents. Still, with regards to the environmental relevance, recorded microplastic concentrations are lower than in our study, and environmental zooplankton samples contain 20 times to four orders of magnitude less PCBs than the post-depuration daphnids [37–39]. Therefore, any current microplastic effects on elimination rates in zooplankton would be small.

Our experiment was designed to test whether depuration via microplastics may occur. Using fish as a model organism, a similar depuration study has been conducted by Rummel and colleagues (2016), who observed no changes in the PCB elimination rate when plastic was added to the fish diet [40]. To the best of our knowledge, an indication of plastic acting as a sink has only been seen in accumulation studies, where POP uptake is reduced in the presence of microplastic [e.g.,17]. While the relative importance of microplastic for bioaccumulation of POPs, compared other matrices in the aquatic environment, is considered small – exploring contamiant transfer is relevant for understandning microplastic toxicity. Future studies should focus on describing both the physiochemical properties of the test microplastic, essential for predicting polymer sorptive capacity [41, 42], and, account for the microplastic effects on test organsisms biochemical composition due e.g., changes in food quality. There is, thereby, still a need of experimental data to quantify POP dynamics in systems containing microplastic for improving our understanding of their role in bioaccumulation of various contaminants, by different organisms, and at different pollution levels [19,42].

Our second hypothesis that lowered PCB body burden in the microplastic-exposed daphnids would result in higher fitness was also partially supported. The addition of microplastic to the food did not affect growth, altough, the reproductive output increased. However, the positive effect of microplastic on the daphnid reproduction was only significant in combination with PCB exposure and coincided with the higher elimination of PCB 209 in the daphnids exposed to microplastic. Overall, the PCB effects observed in the experiment are consistent with the relatively low toxicity of non-dioxin-like PCBs in daphnids [43].

Interestingly, the daphnids fed microplastic-algae mixtures had 1% lower carbon content, while no effect on the %N content was observed, which may indicate subtle effects on energy budget and higher energy allocation to maintenance as well as changes in body stiochiometry. However, without analysis of the biochemical composition, particularly lipid and protein content, this difference is difficult to attribute to a specific physiological change. An increased allocation to reproductive growth is often linked to higher %C content as eggs contain more carbon per dry mass than somatic tissues [44]. However, no individual-level responses, such as fecundity or growth, coincided with the elevated %C content (Fig 3C and D). One can speculate, that the higher fecundity observed in the PCB-challenged daphnids exposed to microplastics led to changes in their metabolism, e.g. more extensive use of lipids [45]. As proteins are relatively low in carbon compared to the lipids, these changes can imply alterations in the elemental composition, such as the decrease in %C levels of somatic tissues and eggs. Moreover, a higher number of embryos in the brood under PCB-induced stress would increase embryonic respiration losses, which may lead to lower %C of the bulk biomass [46].

An alternative explanation to the higher size-specific fecundity in PCB-contaminated daphnids exposed to microplastic is that this occurred as an adaptation to a compromised food quality, because nutrient content per particle was lower in the microplastic algae mixtures. Thus, it is plausible that mixtures with non-palatable particles are perceived by the daphnids as a poor quality food. Poor feeding conditions may lead to a preferential investment in progeny rather than to somatic tissues [47,48]. In this case, a trade-off can manifest as an increased production of smaller offspring, a commonly observed cladoceran response to deteriorated feeding environment [46]. Such responses have been observed in zooplankton that reduced filtering rates at high abundances of inert particles [8,49]. In combination with PCB contamination, these poor feeding conditions may have caused a counterintuitive response manifested as the simultaneous decrease in %C content and the increase in fecundity. Unfortunately, it was not feasible to measure the offspring size, which would have helped to understand the combined effects of microplastics and contaminant exposure on the elemental composition and life histories in this experiment.

The interplay between growth, elemental composition and PCB body burden in the microplastic-exposed daphnids is intriguing and provides insights into the experimental design and informative endpoints to measure. The body burden of hydrophobic chemicals has a positive relation to the organism lipid content, including zooplankton [50]. Effects on the lipid metabolism would, therefore, affect contaminant accumulation, but also growth performance, which may differ ontogenetically as well as among species. Various effects of nanoparticles and microplastics on lipid metabolism have been reported in fish [51] and invertebrates [52,53], with both increased and decreased lipid accumulation. Microplastics have also been found to affect lipid metabolism in zebrafish [54]. Thus, the observed changes in %C content in the daphnids may indicate alterations in lipid metabolism. Such changes suggest that the lipidome profile and biomarkers of lipid synthesis pathways can be of interest in microplastic effect studies, together with the involved physiological (respiration, reproduction, assimilation efficiency and offspring quality) and stoichiometric (C:N and C:P ratios) responses.

## Conclusions

Considering that reported environmental concentrations of microplastics are orders of magnitude lower than those used in our study [55] and that microplastics contribute very little to the ambient concentrations of suspended solids, their *in situ* effects on PCB body burden, growth, and reproduction in zooplankton are most likely negligible. However, the exposure of PCB-contaminated daphnids to microplastic facilitated elimination of the high-molecular-weight PCB by 4-fold compared to the control animals receiving only algal food, which provides proof-of-principle that microplastics can act as a sink for hydrophobic organic contaminants. Although subtle, the decreased carbon content may indicate important metabolic and stoichimetric changes in the animals exposed to microplastic. Given the ecological importance of daphnids (and primary consumers at large) in nutrient recycling, these changes may represent an important mechanism of ecosystem-level responses. Future studies need to focus on the improved mechanistic understanding of the microplastic effects across the biological organization levels.

## Acknowledgements

This study was supported by the Swedish Research Council for Environment, Agricultural Sciences and Spatial Planning (FORMAS, grant number: 216-2013-1010), Isotope Ecology Network in the Baltic Sea region (Swedish Institute, Sweden) (NSSIA project, grant number 12906/2013), the Joint Programming Initiative Healthy and Productive Seas and Oceans (JPI Oceans, WEATHER-MIC project, grant number 942-2015-1866) and the Swedish Innovation Agency VINNOVA and the joint Baltic Sea research and development programme (BONUS, Blue Baltic) for MICROPOLL project [grant number 2017-00979].

## Supporting information

S1 Text: Establishing PCB exposure concentrations S2 Text: Mass-balance calculations for PCBs in the test system. S3 Text: Quality Assurance for PCB analysis S4 Text: Quality Assurance for elemental composition

## References

1. GESAMP. Sources, fate and effects of microplastics in the marine environment: a global assessment. 2015 p. 96. Report No.: 90.

2. Andrady AL. Microplastics in the marine environment. Mar Pollut Bull. 2011;62: 1596–1605. doi:10.1016/j.marpolbul.2011.05.030

3. Cole M, Lindeque P, Halsband C, Galloway TS. Microplastics as contaminants in the marine environment: A review. Mar Pollut Bull. 2011;62: 2588–2597. doi:10.1016/j.marpolbul.2011.09.025

4. Boerger CM, Lattin GL, Moore SL, Moore CJ. Plastic ingestion by planktivorous fishes in the North Pacific Central Gyre. Mar Pollut Bull. 2010;60: 2275–2278. doi:10.1016/j.marpolbul.2010.08.007

5. Browne MA, Dissanayake A, Galloway TS, Lowe DM, Thompson RC. Ingested Microscopic Plastic Translocates to the Circulatory System of the Mussel, *Mytilus edulis* (L.). Environ Sci Technol. 2008;42: 5026–5031. doi:10.1021/es800249a

6. Thompson RC, Olsen Y, Mitchell RP, Davis A, Rowland SJ, John AWG, et al. Lost at Sea: Where Is All the Plastic? Science. 2004;304: 838–838. doi:10.1126/science.1094559

7. Cole M, Lindeque P, Fileman E, Halsband C, Goodhead R, Moger J, et al. Microplastic Ingestion by Zooplankton. Environ Sci Technol. 2013;47: 6646–6655. doi:10.1021/es400663f

8. Cole M, Lindeque P, Fileman E, Halsband C, Galloway TS. The Impact of Polystyrene Microplastics on Feeding, Function and Fecundity in the Marine Copepod *Calanus helgolandicus*. Environ Sci Technol. 2015;49: 1130–1137. doi:10.1021/es504525u

9. Bejgarn S, MacLeod M, Bogdal C, Breitholtz M. Toxicity of leachate from weathering plastics: An exploratory screening study with *Nitocra spinipes*. Chemosphere. 2015;132: 114–119. doi:10.1016/j.chemosphere.2015.03.010

10. Rochman CM, Kurobe T, Flores I, Teh SJ. Early warning signs of endocrine disruption in adult fish from the ingestion of polyethylene with and without sorbed chemical pollutants from the marine environment. Sci Total Environ. 2014;493: 656–661. doi:10.1016/j.scitotenv.2014.06.051

11. Besseling E, Wang B, Lürling M, Koelmans AA. Nanoplastic Affects Growth of *S. obliquus* and Reproduction of *D. magna*. Environ Sci Technol. 2014;48: 12336–12343. doi:10.1021/es503001d

12. Rochman CM, Hoh E, Kurobe T, Teh SJ. Ingested plastic transfers hazardous chemicals to fish and induces hepatic stress. Sci Rep. 2013;3. doi:10.1038/srep03263

13. Brennecke D, Duarte B, Paiva F, Caçador I, Canning-Clode J. Microplastics as vector for heavy metal contamination from the marine environment. Estuar Coast Shelf Sci. 2016; doi:10.1016/j.ecss.2015.12.003

14. Teuten EL, Rowland SJ, Galloway TS, Thompson RC. Potential for Plastics to Transport Hydrophobic Contaminants. Environ Sci Technol. 2007;41: 7759–7764. doi:10.1021/es071737s

15. Velzeboer I, Kwadijk CJAF, Koelmans AA. Strong Sorption of PCBs to Nanoplastics, Microplastics, Carbon Nanotubes, and Fullerenes. Environ Sci Technol. 2014;48: 4869–4876. doi:10.1021/es405721v

16. Avio CG, Gorbi S, Milan M, Benedetti M, Fattorini D, d’Errico G, et al. Pollutants bioavailability and toxicological risk from microplastics to marine mussels. Environ Pollut. 2015;198: 211–222. doi:10.1016/j.envpol.2014.12.021

17. Chua EM, Shimeta J, Nugegoda D, Morrison PD, Clarke BO. Assimilation of Polybrominated Diphenyl Ethers from Microplastics by the Marine Amphipod, *Allorchestes Compressa*. Environ Sci Technol. 2014;48: 8127–8134. doi:10.1021/es405717z

18. Syberg K, Nielsen A, Khan FR, Banta GT, Palmqvist A, Jepsen PM. Microplastic potentiates triclosan toxicity to the marine copepod *Acartia tonsa* (Dana). J Toxicol Environ Health A. 2017;80: 1369–1371. doi:10.1080/15287394.2017.1385046

19. Ziccardi LM, Edgington A, Hentz K, Kulacki KJ, Kane Driscoll S. Microplastics as vectors for bioaccumulation of hydrophobic organic chemicals in the marine environment: A state-of-the-science review. Environ Toxicol Chem. 2016;35: 1667–1676. doi:10.1002/etc.3461

20. Besseling E, Foekema EM, van den Heuvel-Greve MJ, Koelmans AA. The Effect of Microplastic on the Uptake of Chemicals by the Lugworm *Arenicola marina* (L.) under Environmentally Relevant Exposure Conditions. Environ Sci Technol. 2017;51: 8795–8804. doi:10.1021/acs.est.7b02286

21. Bakir A, O’Connor IA, Rowland SJ, Hendriks AJ, Thompson RC. Relative importance of microplastics as a pathway for the transfer of hydrophobic organic chemicals to marine life. Environ Pollut. 2016;219: 56–65. doi:10.1016/j.envpol.2016.09.046

22. Koelmans AA, Besseling E, Wegner A, Foekema EM. Plastic as a Carrier of POPs to Aquatic Organisms: A Model Analysis. Environ Sci Technol. 2013;47: 7812–7820. doi:10.1021/es401169n

23. Koelmans AA. Modeling the Role of Microplastics in Bioaccumulation of Organic Chemicals to Marine Aquatic Organisms. A Critical Review. In: Bergmann M, Gutow L, Klages M, editors. Marine Anthropogenic Litter. Springer International Publishing; 2015. pp. 309–324. doi:10.1007/978-3-319-16510-3_11

24. Nybom I, Abel S, Waissi G, Väänänen K, Mäenpää K, Leppänen MT, et al. Effects of Activated Carbon on PCB Bioaccumulation and Biological Responses of *Chironomus riparius* in Full Life Cycle Test. Environ Sci Technol. 2016;50: 5252–5260. doi:10.1021/acs.est.6b00991

25. OECD. Test No. 202: *Daphnia* sp. Acute Immobilisation Test [Internet]. Paris: Organisation for Economic Co-operation and Development; 2004. Available: http://www.oecd-ilibrary.org/content/book/9789264069947-en

26. Demkowicz-Dobrzański K, Nalecz-Jawecki G, Wawer Z, Bargieł T, Zak M, Sawicki J. Toxicity of neutralized wastes following a synthesis of chlorinated hydrocarbons to *Daphnia magna* and *Chlorella* sp. Sci Total Environ. 1993;134: 1151–1158. doi:10.1016/S0048-9697(05)80119-6

27. Ogonowski M, Schür C, Jarsén Å, Gorokhova E. The Effects of Natural and Anthropogenic Microparticles on Individual Fitness in *Daphnia magna*. PLOS ONE. 2016;11: e0155063. doi:10.1371/journal.pone.0155063

28. Furuhagen S, Liewenborg B, Breitholtz M, Gorokhova E. Feeding Activity and Xenobiotics Modulate Oxidative Status in *Daphnia magna*: Implications for Ecotoxicological Testing. Environ Sci Technol. 2014;48: 12886–12892. doi:10.1021/es5044722

29. Dumont HJ, Velde IV de, Dumont S. The dry weight estimate of biomass in a selection of *Cladocera*, *Copepoda* and *Rotifera* from the plankton, periphyton and benthos of continental waters. Oecologia. 1975;19: 75–97. doi:10.1007/BF00377592

30. Hawker DW, Connell DW. Octanol-water partition coefficients of polychlorinated biphenyl congeners. Environ Sci Technol. 1988;22: 382–387.

31. Herzke D, Anker-Nilssen T, Nøst TH, Götsch A, Christensen-Dalsgaard S, Langset M, et al. Negligible Impact of Ingested Microplastics on Tissue Concentrations of Persistent Organic Pollutants in Northern Fulmars off Coastal Norway. Environ Sci Technol. 2016;50: 1924–1933. doi:10.1021/acs.est.5b04663

32. Goerke H, Weber K. Species-specific elimination of polychlorinated biphenyls in estuarine animals and its impact on residue patterns. Mar Environ Res. 2001;51: 131–149. doi:10.1016/S0141-1136(00)00036-2

33. Grilo TF, Cardoso PG, Pato P, Duarte AC, Pardal MA. Uptake and depuration of PCB-153 in edible shrimp *Palaemonetes varians* and human health risk assessment. Ecotoxicol Environ Saf. 2014;101: 97–102. doi:10.1016/j.ecoenv.2013.12.020

34. Landrum PF, Tigue EA, Driscoll SK, Gossiaux DC, Van Hoof PL, Gedeon ML, et al. Bioaccumulation of PCB Congeners by *Diporeia* spp.: Kinetics and Factors Affecting Bioavailability. J Gt Lakes Res. 2001;27: 117–133. doi:10.1016/S0380-1330(01)70627-2

35. Ge J, Woodward LA, Li QX, Wang J. Distribution, Sources and Risk Assessment of Polychlorinated Biphenyls in Soils from the Midway Atoll, North Pacific Ocean. PLOS ONE. 2013;8: e71521. doi:10.1371/journal.pone.0071521

36. McManus GB, Wyman KD, Peterson WT, Wurster CF. Factors affecting the elimination of PCBs in the marine copepod *Acartia tonsa*. Estuar Coast Shelf Sci. 1983;17: 421–430. doi:10.1016/0272-7714(83)90127-0

37. Tiano M, Tronczyński J, Harmelin-Vivien M, Tixier C, Carlotti F. PCB concentrations in plankton size classes, a temporal study in Marseille Bay, Western Mediterranean Sea. Mar Pollut Bull. 2014;89: 331–339. doi:10.1016/j.marpolbul.2014.09.040

38. Frouin H, Dangerfield N, Macdonald RW, Galbraith M, Crewe N, Shaw P, et al. Partitioning and bioaccumulation of PCBs and PBDEs in marine plankton from the Strait of Georgia, British Columbia, Canada. Prog Oceanogr. 2013;115: 65–75. doi:10.1016/j.pocean.2013.05.023

39. Peltonen H, Ruokojärvi P, Korhonen M, Kiviranta H, Flinkman J, Verta M. PCDD/Fs, PCBs and PBDEs in zooplankton in the Baltic Sea—Spatial and temporal shifts in the congener-specific concentrations. Chemosphere. 2014;114: 172–180. doi:10.1016/j.chemosphere.2014.04.026

40. Rummel CD, Adolfsson-Erici M, Jahnke A, MacLeod M. No measurable “cleaning” of polychlorinated biphenyls from Rainbow Trout in a 9 week depuration study with dietary exposure to 40% polyethylene microspheres. Environ Sci: Processes Impacts. 2016;18: 788–795. doi:10.1039/C6EM00234J

41. Seidensticker S, Zarfl C, Cirpka OA, Fellenberg G, Grathwohl P. Shift in Mass Transfer of Wastewater Contaminants from Microplastics in the Presence of Dissolved Substances. Environ Sci Technol. 2017;51: 12254–12263. doi:10.1021/acs.est.7b02664

42. Hartmann NB, Rist S, Bodin J, Jensen LH, Schmidt SN, Mayer P, et al. Microplastics as vectors for environmental contaminants: Exploring sorption, desorption, and transfer to biota. Integr Environ Assess Manag. 2017;13: 488–493. doi:10.1002/ieam.1904

43. Dillon TM, Benson WH, Stackhouse RA, Crider AM. Effects of selected PCB congeners on survival, growth, and reproduction in *Daphnia magna*. Environ Toxicol Chem. 1990;9: 1317–1326. doi:10.1002/etc.5620091013

44. Sperfeld E, Wacker A. Effects of temperature and dietary sterol availability on growth and cholesterol allocation of the aquatic keystone species *Daphnia*. J Exp Biol. 2009;212: 3051–3059. doi:10.1242/jeb.031401

45. Filho TUB, Soares AMVM, Loureiro S. Energy budget in *Daphnia magna* exposed to natural stressors. Environ Sci Pollut Res. 2010;18: 655–662. doi:10.1007/s11356-010-0413-0

46. Glazier DS. Separating the respiration rates of embryos and brooding females of Daphnia magna: implications for the cost of brooding and the allometry of metabolic rate. Limnol Oceanogr. 1991;36: 354–361.

47. Sibly RM, Calow P. A life-cycle theory of responses to stress. Biol J Linn Soc. 1989;37: 101–116.

48. Stearns SC. Trade-Offs in Life-History Evolution. Funct Ecol. 1989;3: 259–268. doi:10.2307/2389364

49. Kirk KL. Suspended clay reduces *Daphnia* feeding rate. Freshw Biol. 1991;25: 357–365. doi:10.1111/j.1365-2427.1991.tb00498.x

50. Fisk AT, Stern GA, Hobson KA, Strachan WJ, Loewen MD, Norstrom RJ. Persistent Organic Pollutants (POPs) in a Small, Herbivorous, Arctic Marine Zooplankton (*Calanus hyperboreus*): Trends from April to July and the Influence of Lipids and Trophic Transfer. Mar Pollut Bull. 2001;43: 93–101. doi:10.1016/S0025-326X(01)00038-8

51. Cedervall T, Hansson L-A, Lard M, Frohm B, Linse S. Food Chain Transport of Nanoparticles Affects Behaviour and Fat Metabolism in Fish. PLOS ONE. 2012;7: e32254. doi:10.1371/journal.pone.0032254

52. Gomes SIL, Scott-Fordsmand JJ, Amorim MJB. Cellular Energy Allocation to Assess the Impact of Nanomaterials on Soil Invertebrates (Enchytraeids): The Effect of Cu and Ag. Int J Environ Res Public Health. 2015;12: 6858–6878. doi:10.3390/ijerph120606858

53. Wright SL, Rowe D, Thompson RC, Galloway TS. Microplastic ingestion decreases energy reserves in marine worms. Curr Biol. 2013;23: R1031–R1033. doi:10.1016/j.cub.2013.10.068

54. Lu Y, Zhang Y, Deng Y, Jiang W, Zhao Y, Geng J, et al. Uptake and Accumulation of Polystyrene Microplastics in Zebrafish (*Danio rerio*) and Toxic Effects in Liver. Environ Sci Technol. 2016;50: 4054–4060. doi:10.1021/acs.est.6b00183

55. Phuong NN, Zalouk-Vergnoux A, Poirier L, Kamari A, Châtel A, Mouneyrac C, et al. Is there any consistency between the microplastics found in the field and those used in laboratory experiments? Environ Pollut. 2016;211: 111–123. doi:10.1016/j.envpol.2015.12.035

